# Comparative analysis of amphibian genomes: an emerging resource for basic and applied research

**DOI:** 10.1101/2023.02.27.530355

**Authors:** Tiffany A. Kosch, Andrew J. Crawford, Rachel Lockridge Mueller, Katharina C. Wollenberg Valero, Megan L. Power, Ariel Rodríguez, Lauren A. O’Connell, Neil D. Young, Lee F. Skerratt

**Affiliations:** Faculty of Science, University of Melbourne, Melbourne, Australia; Departamento de Ciencias Biológicas, Universidad de los Andes, Bogotá, Colombia; Department of Biology, Colorado State University, Colorado, USA; School of Biology and Environmental Science, University College Dublin, Dublin, Ireland; Institute of Zoology, University of Veterinary Medicine of Hannover, Hannover, Germany; Department of Biology, Stanford University, California, USA

**Keywords:** Amphibian genomes, comparative genomics, transposable elements, repeat expansion, genome synteny

## Abstract

Amphibians are the most threatened group of vertebrates and are in dire need of conservation intervention to ensure their continued survival. They exhibit unique features including a high diversity of reproductive strategies, permeable and specialized skin capable of producing toxins and antimicrobial compounds, multiple genetic mechanisms of sex determination, and in some lineages, the ability to regenerate limbs and organs. Although genomics approaches would shed light on these unique traits and aid conservation, sequencing and assembly of amphibian genomes has lagged behind other taxa due to their comparatively large genome sizes. Fortunately, the development of long-read sequencing technologies and initiatives has led to a recent burst of new amphibian genome assemblies. Although growing, the field of amphibian genomics suffers from the lack of annotation resources, tools for working with challenging genomes, and lack of high-quality assemblies in multiple clades of amphibians. Here we analyze 51 publicly available amphibian genomes to evaluate their usefulness for functional genomics research. We report considerable variation in genome assembly quality and completeness, and report some of the highest transposable element and repeat contents of any vertebrate. Additionally, we detected an association between transposable element content and climatic variables. Our analysis provides evidence of conserved genome synteny despite the long divergence times of this group, but we also highlight inconsistencies in chromosome naming and orientation across genome assemblies. We discuss sequencing gaps in the phylogeny and suggest key targets for future sequencing endeavors. Finally, we propose increased investment in amphibian genomics research to promote their conservation.

## INTRODUCTION

Amphibians are an ancient lineage of vertebrates that predate amniotes by more than 100 million years. Despite the considerable age of this lineage, amphibians are now the most threatened group of vertebrates with more that 40% of species and are threatened by factors such as habitat change, disease, and over-exploitation (IUCN, 2022; Scheele et al., 2019). Notably, many of these threats are hard to reverse, suggesting that novel approaches that utilize genomic resources may lead to improved management decisions for some of the most endangered taxa (Kosch et al., 2022; Scheele et al., 2014).

We are only just beginning to understand the genetic basis of many of the unique features of amphibians. Amphibians exhibit a high diversity of reproductive strategies including biphasic and direct development, uniparental and biparental care, mouth and gastric brooding, and foam nesting (Brown et al., 2010; Nunes-de-Almeida et al., 2021; Schulte et al., 2020). They also have specialized skin capable of producing complex compounds of interest for drug discovery for the development of antimicrobial drugs and analgesics (Daly et al., 2000; De Angelis et al., 2021; Liu et al., 2020).

Amphibians occur across habitat types from rainforests to deserts, freshwater streams to salt marshes, and tropical to arctic climates (Duellman, 1999), but it is unclear how this ecological diversity is reflected in genome composition. One potential way is the number of transposable elements (TEs) present in the genome. TEs have a huge impact on the structure and function of eukaryotic genomes, with amphibians having among the largest TE content among vertebrates. There is increasing evidence that TE activity, and thus their relative proportion in genomes, is influenced by abiotic factors (Pimpinelli & Piacentini, 2020). This in turn highlights their potential role in the regulation of genetic mechanisms responsible for environmental adaptation (Casacuberta & González, 2013; Pappalardo et al., 2021). Salamanders are an important resource for transplant and regeneration research due to their ability to regenerate limbs and internal organs (Elewa et al., 2017; Nowoshilow et al., 2018). Amphibians also have many of the same immune components of mammals making them an important model resource for immunology (Paiola et al., 2023; Robert, 2020).

Despite the obvious value of amphibian genomes for research on ecology, evolution, medicine, and improving their conservation, until recently, the generation of amphibian reference genomes has been markedly slower than other vertebrates (Hotaling et al., 2021a; Womack et al., 2022). This lag can be attributed to high costs and the computational challenges of assembling their often large and complex genomes (Sun et al., 2020). Recent advances in sequencing technologies such as long read sequencing and assembly algorithms that incorporate hybrid approaches have circumvented many of these challenges leading to a surge of high quality, chromosome-level reference genomes. The next challenge will be developing the tools for annotation and comparative analyses of these large genomes.

In this study, we provide a synthesis of all available amphibian reference genome assemblies, 51 at the time of our analysis, with the number growing every day. We evaluate assembly quality, sequencing technology, gene completeness, transposable element and repeat content and its ecological correlates, taxonomic representation, and synteny.

## MATERIALS AND METHODS

### Genomes

A search of the NCBI genome website using the search term “amphibians” conducted on August 25, 2023, revealed there were 90 amphibian genomes from 68 species. All genome files in fasta format were downloaded for assessment. Sixteen salamander genomes (Pyron et al., 2024) were excluded from our analyses due to their high degree of incompleteness (i.e., <10% of genome assembled). Of the remaining genomes, one genome was selected for each species for subsequent analysis. If there was more than one draft of a genome, the most recent draft and/or the primary haplotype was selected. In cases where there were multiple versions sequenced by different groups, the best genome was selected by lowest scaffold number. Entire genomes (including uncharacterized contigs but excluding mitochondrial genomes) were used for assessment unless indicated otherwise.

Genome databases NCBI Genomes, NCBI RefSeq (O’Leary et al., 2016), Ensembl (Cunningham et al., 2022), UCSC Genome Browser (Lee et al., 2022), and Genomes on a Tree (GoaT) (Sotero-Caio et al., 2021) were searched for information on the 51 amphibian species with reference genomes including chromosome number, annotation data, proteome availability, C-value, and sequencing technology. Sequencing strategy was classified as “short-single” for Illumina only sequencing, “long-single” for sequencing using long read technologies (e.g., PacBio and Oxford Nanopore), and “hybrid” for sequencing approaches using more than one approach (e.g., PacBio and Hi-C).

A search for amphibian proteome datasets on NCBI RefSeq (O’Leary et al., 2016), Ensembl (Cunningham et al., 2022), and UCSC Genome Browser (Lee et al., 2022) databases on June 24, 2022 revealed 11 proteomics datasets.

A search of the NCBI Organelle database on 15, February 2023 using search term “amphibian” resulted in 353 mitochondrial genomes belonging to 345 species (Table S11). Seventeen mitochondrial genomes overlapped with the amphibian nuclear genomes analyzed in this study.

### Reference genome availability summary

The GoaT online database (Sotero-Caio et al., 2021) was searched on August 28, 2023 to summarize genomes in progress or publicly available using the search terms “tax_tree(Amphibia) AND tax_rank(species) AND sequencing_status=in_progress” or “tax_tree(Amphibia) AND tax_rank(species) AND sequencing_status=insdc_open”. The same search terms were used to summarize publicly available genomes for mammals, birds, and non-avian reptiles with the “tax_tree” search term replaced by appropriate Class.

### Genome quality analyses

Genome quality assessment was performed with BBMap (v. 39.01) “statswrapper.sh” bash script (https://github.com/BioInfoTools/BBMap). This tool generates metrics such as genome size, contig N50, and scaffold count. Benchmarking Universal Single-Copy Orthologs (BUSCO) were summarized with the BUSCO tool (v. 5.1.2) (Manni et al., 2021) using the OrthoDB Tetrapoda ortholog library (v. odb10) (Kriventseva et al., 2018) (N=5310 orthologs) with the prompt “-m genome”. Percentage of the genome assembled to chromosomes was calculated with a custom bash script that computes the genome length assigned to chromosomes and divides it by the “assembly length” value computed by BBMap.

### Phylogenetic tree

A species to family correspondence table was obtained from Jetz and Pyron (2018) (https://vertlife.org/files_20170703/) and was filtered to include only the species with the longest nucleotide sequence per family. This taxa subset was used to obtain a subset of 100 phylogenetic trees from the posterior distribution of the Jetz and Pyron (2018) dataset, as available from http://vertlife.org/phylosubsets. A consensus tree from these 100 trees was then obtained using treeannotator (v2.7.5) (settings - target tree type: maximum clade credibility, node heights: median burn-in percentage: 0, posterior probability limit: 0.0) (Drummond & Rambaut, 2007). The species names of the tree tips were then substituted with the corresponding family names using the “sub.taxa.label” function in the phylotools package (https://github.com/helixcn/phylotools) in R with the aid of the species to family correspondence table, which was updated with the most recent classification available in AmphibiaWeb (https://amphibiaweb.org) and the Amphibian Species of the World database (https://amphibiansoftheworld.amnh.org/). In cases where these two references disagreed, the AmphibiaWeb taxonomy was used.

### Repeat modelling and annotation

Repeats were de novo modelled with RepeatModeler (Apptainer v. 1.2.3) (Flynn et al., 2020). Genomes were then annotated using RepeatMasker (v. 4.1.2-p1) (Smit et al., 2013) with a concatenated library of genome-specific repeats generated from RepeatModeler and the Dfam amphibian repeat library (v. Dfam.h5) (Storer et al., 2021). Before annotation, any previous soft masking of the genomes was reversed. The results were summarized using a custom bash and R scripts.

### Ecological correlates of transposable element content

Occurrence data for the 51 species were downloaded from the Global Biodiversity Information Facility (GBIF) (https://www.gbif/org/; last accessed February 2024 (full DOI’s for each occurrence data set in Table S5). In addition, due to the putative involvement of temperature in TE activity, BioClim variables associated with temperature (Bio1-Bio11) were obtained for the 51 amphibian species (Table S6). As previous studies have explored the relationship between amphibian genome size and environmental variables (Liedtke et al., 2022), here we focused on the relationship between temperature variables, elevation, and amphibian transposable elements. Influence of these bioclimatic variables (after removing highly collinear variables, see supplementary methods) on transposable element content (summarized into three groups: proportion of total transposable elements (TEs), proportion of retroelements, and proportion of DNA transposons) was modelled using Bayesian mixed effect models (Hadfield, 2010). To correct for body size, log transformed body size was included in the model structure with log transformed Bio2 (Mean diurnal range), Bio4 (temperature seasonality), Bio8 (mean temperature of wettest quarter), Bio10 (mean temperature of warmest quarter) and elevation. Models were also corrected for phylogenetic non-independence (Figure S5, see Supplementary Methods for further information) with phylogenetically independent contrasts (Felsenstein, 1985; Garland Jr et al., 1992) plotted with and (Revell, 2024).

### Synteny analysis

Synteny of BUSCO genes for chromosome level assemblies was analyzed with R Package GENESPACE (v. 1.1.4) (Lovell et al., 2022), which uses OrthoFinder (v. 2.5.4) (Emms & Kelly, 2019) to infer orthology. Synteny was analyzed using BUSCO “full_table.tsv” results files that were reformatted for GENESPACE input using a custom bash script. Synteny plots were generated for all chromosome level assemblies, all anuran chromosome level assemblies, for the two salamander genomes, and for the three caecilian genomes using the GENESPACE plotting tool “plot_riparian”. Chromosomes with reversed orientation compared to the reference genome were inverted to improve visualization.

### Quantification and statistical analysis

Regression analyses, ANOVAs, and Student’s t-tests for comparing genome quality measurements were conducted with the R statistics package (v. 4.1.2) (Team, 2013) in R Studio (v. 2022.02.3) (Team, 2022). Genome quality measures, contigN50, and scaffold count, were log transformed prior to analysis. R-scripts for statistical analysis and plotting are available on GitHub at https://doi.org/10.5281/zenodo.7679280.

## RESULTS

### Genome quality

A total of 51 nuclear amphibian genome assemblies were available for our study and were generated with a variety of sequencing technologies, including Illumina (NextSeq, HiSeq), PacBio (RS11, Sequel), and Oxford Nanopore. Sequenced genomes represented 25 of 73 amphibian families with reference genomes distributed unevenly across the phylogeny (Fig. 1). For example, there are only two salamander genomes representing the 798 extant species, no genomes representing anuran families such as Leiopelmatidae or Hyperoliidae, yet there are seven Ranidae and six Pipidae genomes (Fig. 1).

**Figure 1.**
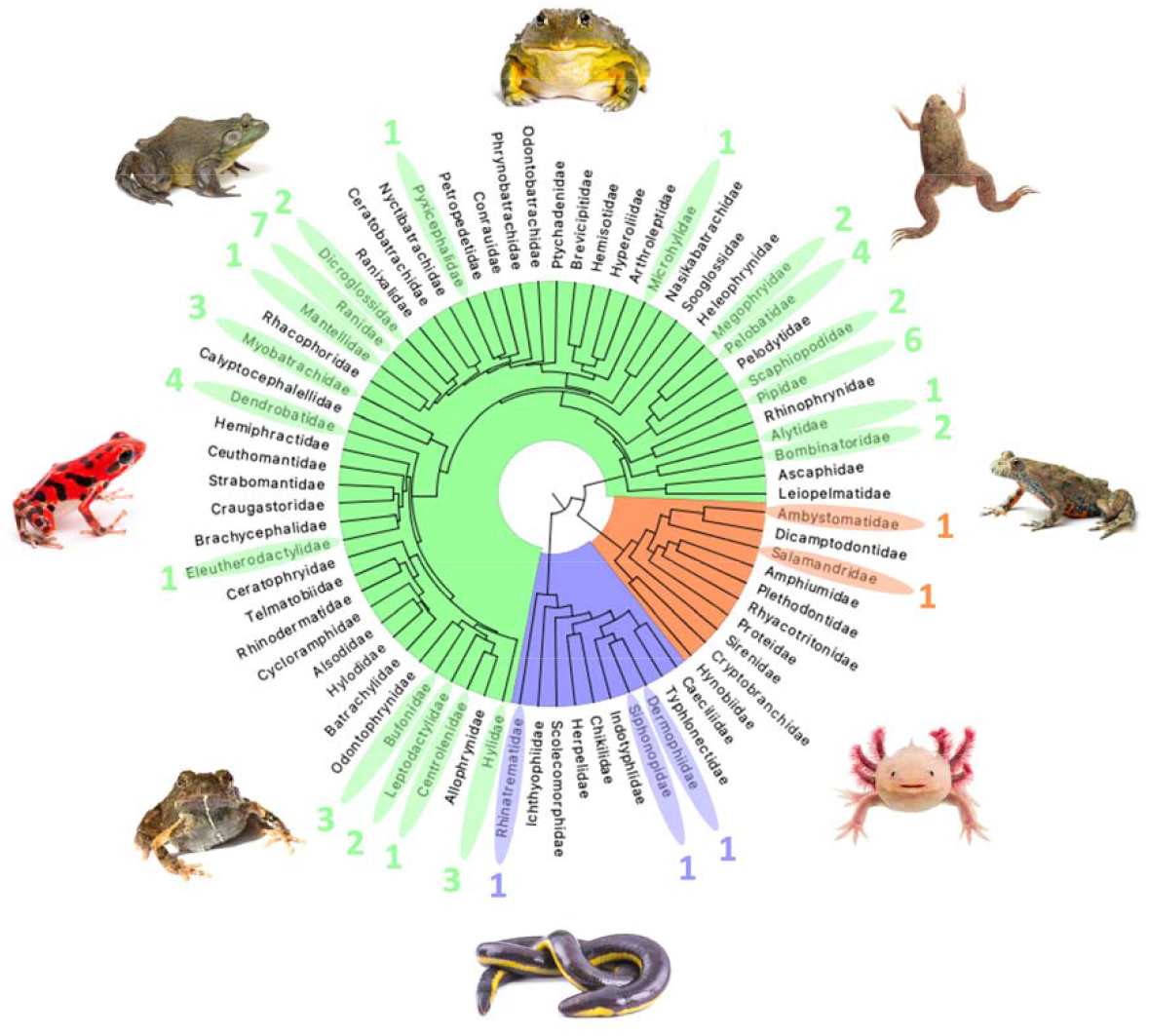
Phylogenetic tree of amphibian families. Amphibian families with representative genomes are highlighted and numbers indicate genome counts per family. (Green) anurans, (blue) caecilians, and (orange) salamanders. *Engystomops pustulosus* (Family) image was taken by B. Gratwicke, other amphibian images were licensed to T. Kosch by Adobe Stock and Shutterstock.

Genome assembly length ranged from 0.48 Gb in *Scaphiopus couchii* to 28.21 Gb in *Ambystoma mexicanum* and was strongly positively associated with c-value estimates of genome size (*F*_49_ = 330.5, p < 1 × 10^-15^) (Table 1, Fig. S1). Twenty-eight of these genomes were assembled to the chromosome level of which the percentage of the genome assigned to chromosomes ranged from 63.88 to 99.99% (Table 1). Percentage of the genome assigned to chromosomes was positively associated with contig N50 (*F*_26_ = 8.6, p = 0.007) and read length (*t*_29.2_ = 3.07, p = 0.005) and negatively associated with the number of scaffolds (*F*_26_ = 25.2, p < 0.00001). There are additionally mitochondrial genome assemblies for 345 species of which 17 had nuclear reference genomes. Eleven of the species with genomes had proteomics data (Table S1).

**Table 1.**
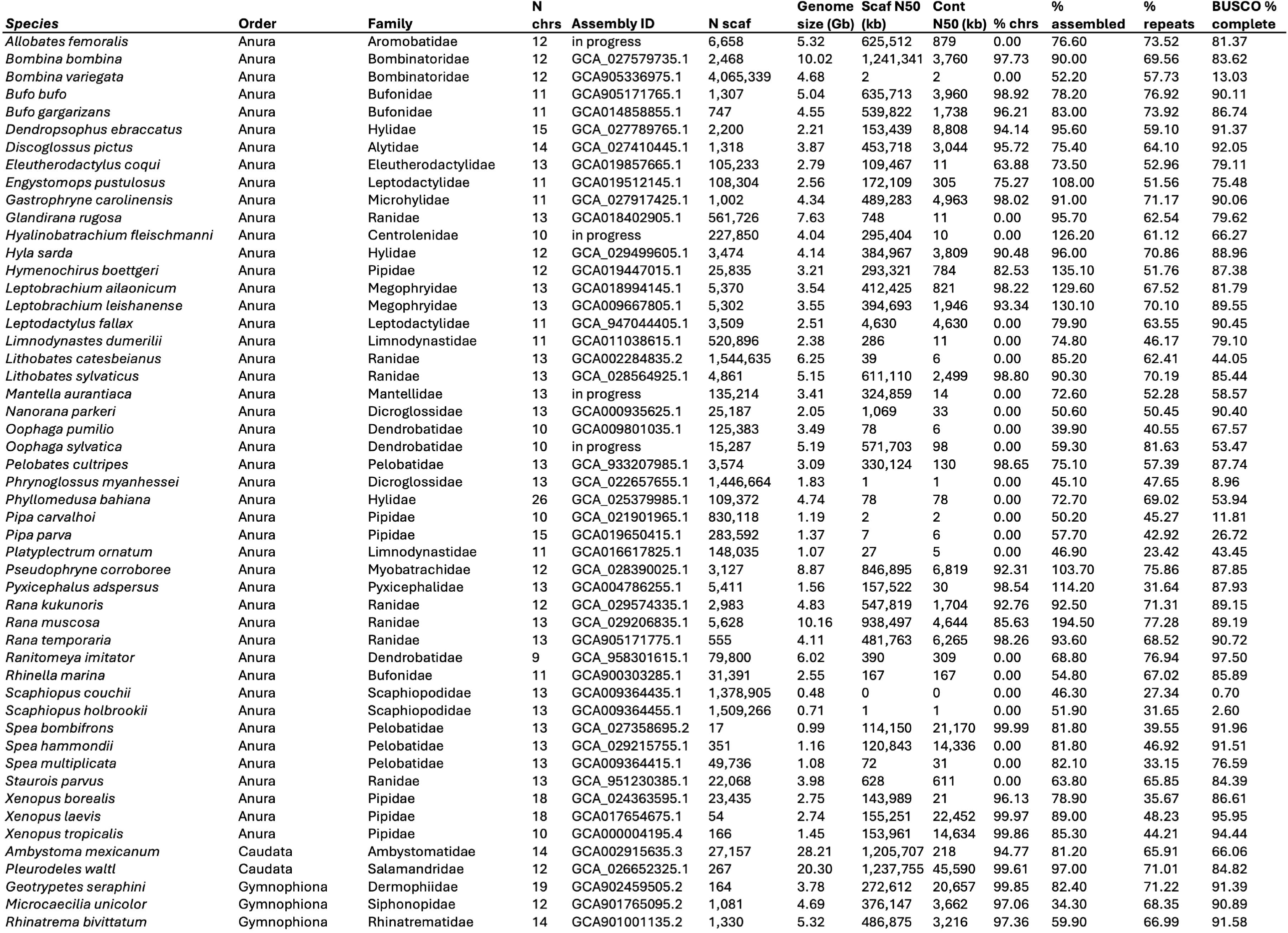
Genome quality measures.

The quality of the amphibian genomes varied considerably (Table 1). Genomes generated with short-read technologies were of lower quality than long-read or hybrid genome assemblies as indicated by significantly lower contig N50s (*F*_2,48_ = 26.91, p < 10^-6^), percentage of complete Benchmarking Universal Single-Copy Ortholog (BUSCO) genes (Fig. S3; *F*_2,48_ = 10.52, p < 0.001), and higher scaffold numbers (*F*_2,48_ = 15.8, p < 10^-5^).

Contig N50 ranged from 362 bp in *S. couchii* to 45.59 Mb in *Pleurodeles waltl* with a median of 611.23 Kb. Scaffold count varied considerably from 17 in *Spea bombifrons* to more than four million in *Bombina variegata* with a median of 6.66 Kb (Table 1). Benchmarking Universal Single-Copy Orthologs (BUSCO) scores ranged from 0.7 to 99.5% completeness (Tables 1, S1; Fig. 2) and were positively associated with contig N50 (F_49_ = 82.6, p < 10-^10^; Fig. S2) and scaffold count (F_49_ = 66.04, p < 10^-8^). Most genomes had low percentages of duplicate BUSCO genes (< 6%), suggesting they may be diploid except for *Ranitomeya imitator* and the known tetraploid species, *X. laevis and X. borealis* (Fig. 2) (Tymowska & Fischberg, 1973).

**Figure 2.**
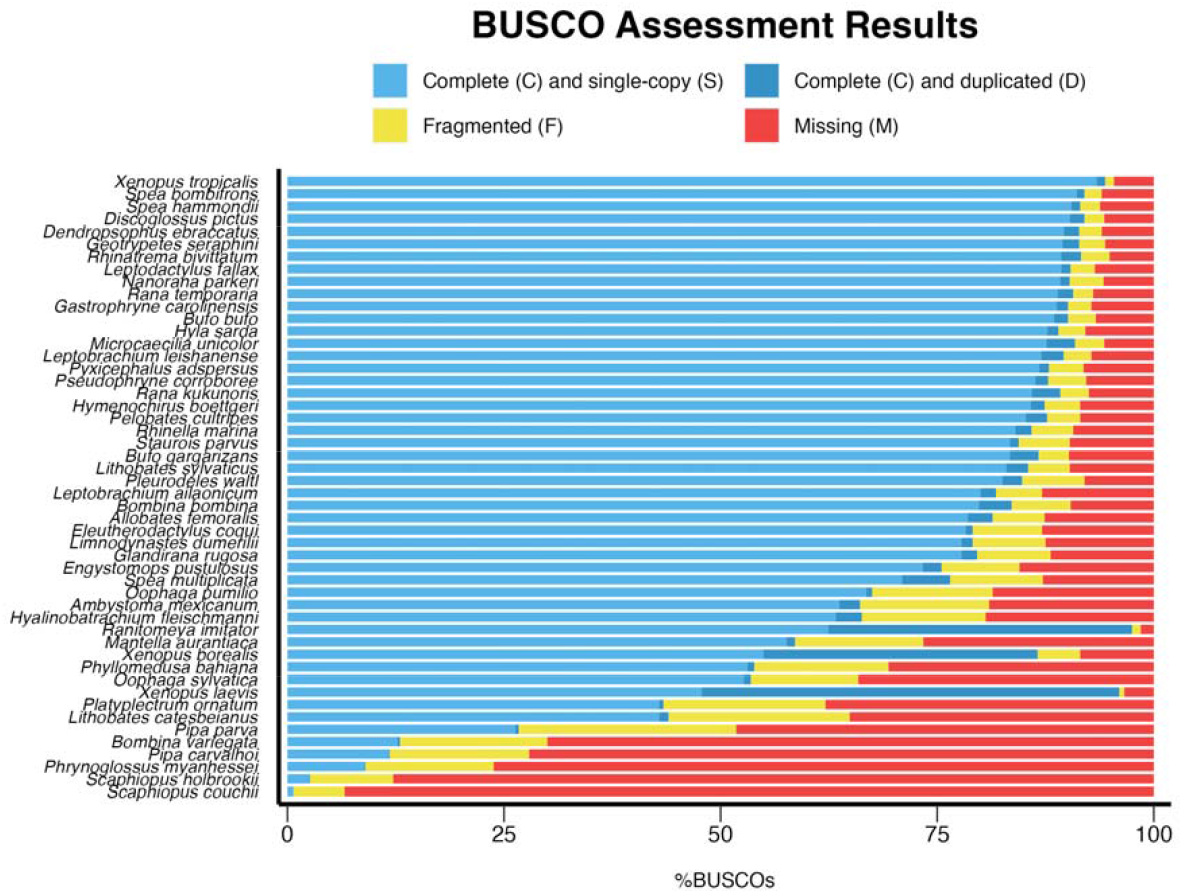
BUSCO (Benchmarking Universal Single-Copy Orthologs) assessment results for amphibian genomes.

### Repeat content

Overall identified repeat percentage of the genomes ranged from 23% in *Platyplectrum ornatum* to 82% in *Oophaga sylvatica* and was positively associated with genome size (F_49_ = 13.24, p = 0.0006) (Tables 1; Fig. S3). Repeat content varied across genomes with the anurans *Pseudophrne corroboree, Bombina bombina*, and *O. sylvatica* dominated by Long Terminal Repeats (LTRs), the three caecilians dominated by Long Interspersed Nuclear Elements (LINEs), and many of the ranid and bufonid anurans dominated by DNA transposons (Fig. 3; Tables S2-S4). Salamander genomes *Ambystoma mexicanum* and *Pleurodeles waltl* had fewer repeats than might be predicted given their large sizes (Fig. S3).

**Figure 3.**
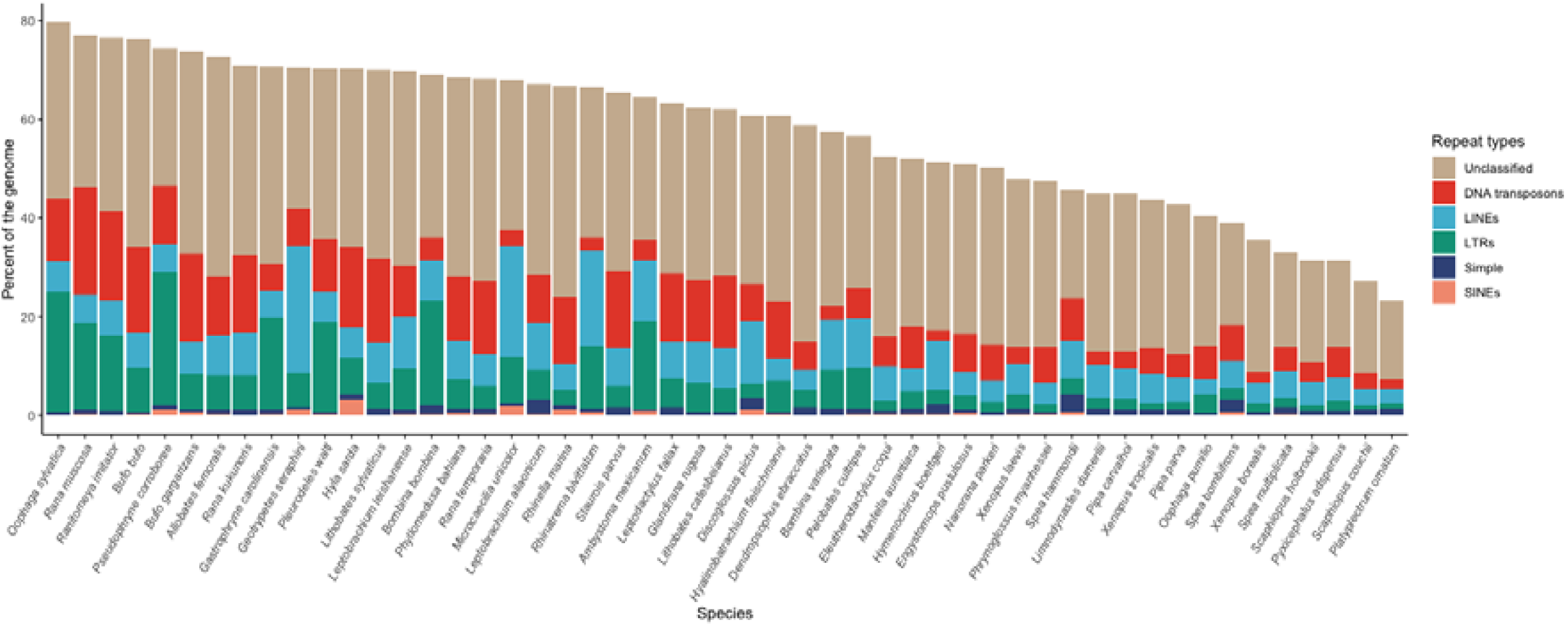
Repeat content across the amphibian genomes. (LINEs) long interspersed nuclear elements, (LTRs) long terminal repeats, and (SINEs) short interspersed nuclear elements.

The proportion of repeats that could be classified by RepeatMasker ranged from 7.4 % in *P. ornatum* to 47.8 % in *P. corroboree* (Table S1) and was positively associated with genome quality measures contigN50 (F_49_ = 23.49, p = 0.001), scaffold count (F_49_ = 8.71, p = 0.005), and percent BUSCO complete (F_49_ = 10.27, p = 0.002). The ability to classify repeats was also positively associated with read length, with longer reads resulting in better classification (t _35.622_ = 4.73, p < 0.001).

### Ecological correlates of transposable element content

A Bayesian mixed effect modelling approach was employed to examine the relationships between proportion of transposable elements and environmental variables. Controlling for phylogenetic relationships (by estimating Pagel’s lambda, λ; de Villemereuil & Nakagawa, 2014), including body size as a covariate (Spearman correlation with transposable element content ⍰= −0.772, p<0.001) and excluding the three globally invasive species (*Rhinella marina, X. laevis*, and *Lithobates catesbeianus*) our analysis revealed a significant influence (pMCMC = 0.014) of Bio8 (mean temperature of the wettest quarter) on the proportion of total transposable elements (Figs 4, S5; Table S8). Inclusion of these three invasive species did not change this relationship (Table S7). Further analysis indicated that the relationship with Bio8 was not specific to a particular class of transposable elements, such as retroelements or DNA transposons (Tables S9 and S10). Phylogenetic signal (Pagel’s lambda, λ) was moderate when considering total transposable elements and retroelements (0.555; Table S7) and increased when we considered retroelements and DNA transposons alone (0.616 and 0.649; Table S9).

**Figure 4.**
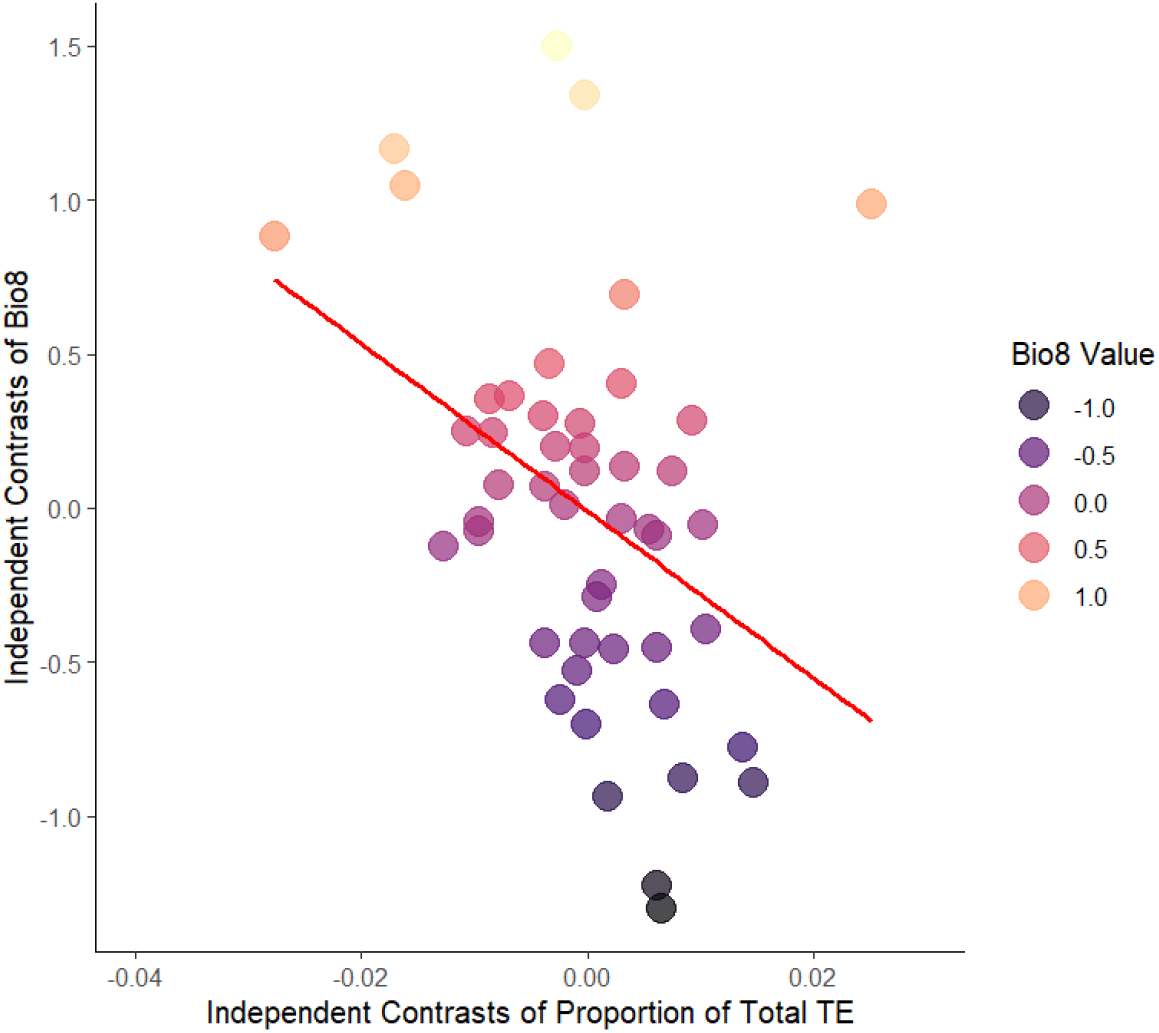
Phylogenetic independent contrasts (PICs) between the proportion of transposable element content relative to genome size and Bio8 (representing mean temperature of wettest quarter).

### Genome synteny

Genome synteny of BUSCO genes was highly conserved within amphibian orders (caecilians (Fig. S7), caudates (Fig. S8), and anurans (Fig. S9); but was less conserved across the amphibian orders (Fig. 5, S6). However, chromosome naming was inconsistent across all taxa (Figs 5, S6-S9). For example, *X. tropicalis* chr1 is chr12 in *Leptobrachium ailaonicum* (but not *L. leishanense*), chr2 in *Bufo bufo* (but not *Bufo gargarizans*) (Fig. S9), and most of the chromosomes for the two salamander genomes (Fig. S8). Orientation of chromosomes was also inconsistent, including between species of the same genus (e.g., *Bufo, Leptobrachium*) (Fig. S59) and among the three caecilians (Fig. S7). Multiple inversions were evident including between chr3 of pipids (*Xenopus tropicalis* and *Hymenochirus boettgeri*) and other anurans (chromosomes 1, 2, 3, 4, or 10), caecilians (chr3 and chr4/5/6), and even within species of the same genus (chr7 *Bufo gargarizans*, chr 9 B. *bufo*; Figs 5, S7, S9). There was also evidence of several chromosomal fissions including the separation of chr1 of *Leptobrachium leishanense* into chr3 and chr6 in *Pyxicephalus adspersus* and into chr3 and chr7 in *Engystomops pustulosus;* however, this chromosome remained mostly intact in the other anuran genomes (Fig. S9).

**Figure 5.**
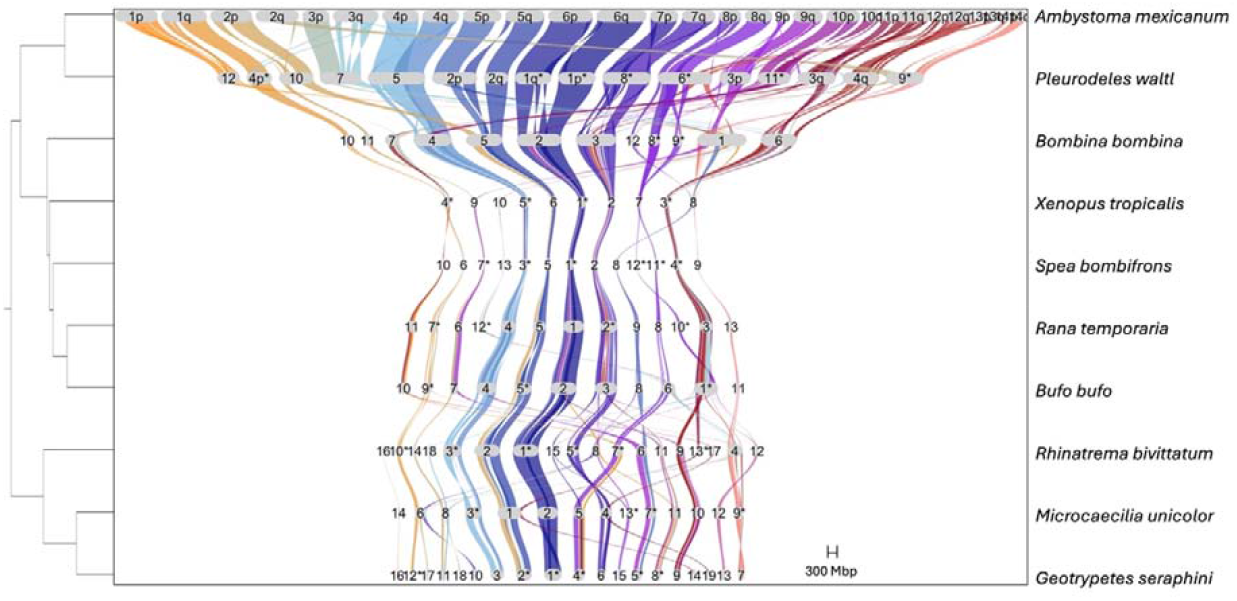
Synteny plot of BUSCOs (Benchmarking Universal Single-Copy Orthologs) for representative amphibian chromosome-level genomes. The phylogenetic tree was created with Timetree.org. The reference genome is *Ambystoma mexicanum*. *Indicate inverted chromosomes. Chromosomes without BUSCOs were excluded from the plot.

## DISCUSSION

In this study we analyzed 51 amphibian reference genomes from the public domain to evaluate their content and usefulness for functional genetics research (Fig. 1, Table 1). There are considerably fewer reference genomes for amphibians than exist for birds (N=754), mammals (N=406), and non-avian reptiles (N=108). This scarcity of reference genomes results in many gaps in genome representation across the amphibian tree of life including many entirely unrepresented groups and with only two genomes representing the entire order Caudata (but see Myers & Pyron, 2024). The unrepresented families include many of interest from a conservation perspective due to their high number of IUCN RedList Critically Endangered species (e.g., Cryptobranchidae, Plethodontidae, Strabomantidae, and Craugastoridae) (IUCN, 2022). However, our search of the Genomes on a Tree (GoaT) database (Sotero-Caio et al., 2021) indicated that there are a further 20 amphibian genome assemblies in progress (15 anurans, 5 caudates; Table S10) indicating that this resource will be increasing by more than 40% in the next few years.

The quality and completeness of the genomes in our dataset varied considerably (e.g., Fig. 2). Much of this variation can be attributed to the sequencing technology used to generate them, with short-read sequencing approaches resulting in lower completeness and continuity (Fig. S2). These impacts are a recognized limitation of short-read sequencing and have been reported to impact genome quality in taxa from insects (Hotaling et al., 2021b) to other vertebrates (Rhie et al., 2021), but have likely had a disproportionate impact in amphibian genomes due to the difficulty of assembling genomes with high repeat content (Sun et al., 2020). Fortunately, most ongoing sequencing efforts now use long-read or hybrid sequencing approaches (i.e., that incorporate scaffolding technologies such as Hi-C sequencing), which along with improved sequencing algorithms, should result in higher quality amphibian genomes (Hotaling et al., 2021a; Lawniczak et al., 2022; Rhie et al., 2021).

The variation we report here in genome quality, contiguity, and completeness may impact the value of the genomes for functional genomics research. However, the improvements in all these measures seen with the utilization of long read technologies or hybrid assemblies suggests that genome quality will continue to improve as these approaches are used more frequently. Genome quality (i.e., high continuity, contiguity, accuracy, completeness (Rhie et al., 2021)) are critical for applications such as quantitative genetics where assembly errors can lead to incorrect inferences in genetic association or genetic prediction. Quality also enhances the usefulness of genomes. For example, highly contiguous chromosome-level assemblies decrease computational requirements for downstream analyses such as mapping, variant calling, and alignment (Aganezov et al., 2022).

One of the most intriguing features of amphibian genomes is the huge range they exhibit in size (Biscotti et al., 2019). This was exemplified in our dataset where assembly length ranged from 0.48 Gb in *Scaphiopus couchii* to 28 Gb in *Ambystoma mexicanum*. Why gigantic genomes exist in some species, but not others, remains a key evolutionary question (Kapusta et al., 2017; Wang et al., 2021). Explanations include differences in genome-level processes (e.g., insertion and deletion rates) (Frahry et al., 2015; Sun et al., 2012b), development (e.g., developmental rate and complexity) (Gregory, 2002; Liedtke et al., 2018), physiology (e.g., water loss) (Johnson et al., 2021), body size (e.g. miniaturization) (Decena-Segarra et al., 2020), and demography (e.g., effective population size) (Liedtke et al., 2018; Lynch & Walsh, 2007) (but see Mohlhenrich & Mueller, 2016). As more amphibian genomes become available, these hypotheses can be more rigorously evaluated.

We report some of the largest estimates of repeat content of any vertebrate (82% in *Oophaga sylvatica* and 77% in *Rana muscosa*), exceeded only by the Australian lungfish at 90% (Meyer et al., 2021). As expected, genome size was correlated with repeat content affirming that much of the variation in amphibian genome size is due to an excess of repeats and transposable elements rather than coding regionds (Biscotti et al., 2019; Lamichhaney et al., 2021; Zuo et al., 2023).

In contrast to mammals, whose repeat landscape is mainly dominated by LTR retrotransposons (Platt et al., 2018), amphibian repeat content varied considerably with some species dominated by DNA transposons (as previously reported (Suda et al., 2022; Zuo et al., 2023), and others by non-LTR retrotransposons including the three caecilian genomes which were dominated by LINEs. This agrees with genomic data and transcriptomic data from the caecilian *Ichthyophis bannanicus*, where LINEs were the second most abundant type of repeat (26% of the genome) behind Dictyostelium intermediate repeat sequences (DIRS) (30%) (Wang et al., 2021); this is a similar percentage of LINES to what we report in the three caecilian genomes in this study (19 to 26%) (Table S4).

These disparities in repeat percentage and content likely reflect differing evolutionary histories among species, as indicated by three of the four congeneric species pairs in our dataset having similar values (i.e., *Bufo, Leptobrachium*, and *Xenopus;* but not *Oophaga*). The differences we observed in *Oophaga pumilio* and *O. sylvatica* are likely due to assembly quality rather than genome content given that these two genomes were sequenced with different technologies and have dramatically different genome qualities (e.g., contig N50s of 5.8 vs. 97.8 Kbp respectively).

A considerable proportion of the repeats could not be classified. This was likely due to incorrect classification (e.g., genes categorized as repeats) and the lack of good amphibian-specific repeat resources (Ou et al., 2019) for classification via nucleotide sequence homology. The majority of amphibian curated repeat libraries are generated in reference to *Xenopus* species (e.g., Dfam); the large divergence times of this genus from the other amphibian species suggests that it may be a contributing factor to the lack of classification. However, we also report many unclassified repeats in the two *Xenopus* genomes.

The largest genomes in our dataset from caudates, *A. mexicanum and P. waltl*, had fewer repeats than predicted given their size (Fig S4) (Nowoshilow et al., 2018). This may be due, in part, to the Dfam (Storer et al., 2021) library used for repeat annotation being anuran-based; however, we did not observe this trend in the three caecilian genomes in our dataset. Also, we performed *de novo* annotation of these genomes, which should have captured repetitive elements missing from Dfam. More likely, this low number of repeats reflects low deletion rates and, thus, persistence of repeats in the genome for extremely long periods of time, leading to their mutational decay into unique sequences whose repetitive origin is obscured (Frahry et al., 2015; Keinath et al., 2015; Novák et al., 2020; Sun et al., 2012a).

We also show that amphibian species that inhabit warm climates particularly during months with high precipitation have a greater proportion of transposable elements. This observed trend does not appear to be driven by a specific group of transposons suggesting it may be caused by climatic factors. Recent studies indicate that transposable elements exhibit greater activity in hotter climates (Baduel et al., 2021) with an increasing number of studies suggesting increased transposable element activity contributes to genetic diversification and facilitates species adaptation (Li et al., 2018; Schrader & Schmitz, 2019; Stapley et al., 2015). The pattern observed here likewise suggests the potential for heightened transposable element activity and may help explain transposable element accumulation and potentially the higher evolutionary rates observed in the genomes of tropical amphibians (Pyron & Wiens, 2013).

Our study is the first to examine chromosomal synteny across all amphibian orders. We show that overall synteny of amphibian genomes is relatively conserved, particularly within orders (Figs 5 and S7). This aligns with previous results from anurans that reported conserved genome organization in this group (Bredeson et al., 2021; Wu et al., 2022). However, chromosome content and number varied across species, which seems to have been driven by multiple occurrences of chromosomal fusions and fissions (e.g., Fig. 5). Chromosomal rearrangements have occurred throughout vertebrate evolution, including the hypothesized fusion of microchromosomes in the ancestor of tetrapods to create the larger macrochromosomes seen in amphibians and mammals and their subsequent fission to create the microchromosomes of modern birds and non-avian reptiles (Waters et al., 2021).

Some of the structural rearrangements we detected may be due to assembly errors and should be evaluated in future assemblies using long-read scaffolding approaches (e.g., Oxford nanopore sequencing), chromosome conformation capture technologies (e.g., Hi-C), or chromosome mapping approaches (e.g., FISH). We also identified incongruities with chromosome naming and orientation caused by differences in assembly methods. These were apparent even within species of the same genus (e.g., *Bufo*). We suggest potential revisions of existing genome annotations to improve congruity and that future assemblies are curated consistently against high-quality reference genomes (e.g., *Xenopus laevis*).

## Conclusions

New sequencing technologies and assembly algorithms have resulted in a good number of genomes for comparative analyses spanning the amphibian phylogeny. This has already begun to yield important insights on the evolution (Lamichhaney et al., 2021; Wu et al., 2022), development (Schloissnig et al., 2021; Stuckert et al., 2021), sex determination (Hime et al., 2019; Ma & Veltsos, 2021), and unique features (Fischer et al., 2019; Nowoshilow et al., 2018; Seidl et al., 2019) of this interesting group of animals.

The increased availability of amphibian genomes can also aid conservation efforts in this highly threatened group by facilitating research on genome-wide functional diversity, which can be used to inform management decisions such as genetic rescue or targeted genetic intervention for species threatened by habitat loss or chytridiomycosis (Chestnut et al., 2014; Kosch et al., 2022). Additionally, well-annotated genomes can be used to create eDNA assays for population monitoring (Breton et al., 2022; Saeed et al., 2022).

Future research efforts should focus on generating more reference genomes to fill the gaps in the amphibian phylogeny and the identification of advantageous genetic traits against threats. Efforts should also be made to increase the quality of genomes and expand transcriptome and annotation databases. We suggest that these efforts strive to follow the recommendations of initiatives such as the Earth BioGenome Project (Lawniczak et al., 2022), the Darwin Tree of Life Project (Blaxter et al., 2022), and the Threatened Species Initiative (Hogg et al., 2022) to sequence at least one representative from each family to ensure taxonomic coverage. Species selection should prioritize species of interest for understanding valuable functional genetic traits; for example, for the purpose of immunological research to understand disease resistance, or for conservation purposes to enhance fitness.

## ACKNOWLEDGEMENTS

T.A.K, N.D.Y, and L.F.S research were supported by The University of Melbourne’s Research Computing Services and the Petascale Campus Initiative. T.A.K. and L.F.S were supported by Australian Research Council Grants (FT190100462, LP200301370). K.W.V and M.L.P were funded by the European Union (ERC, MolStressH2O, 101044202). Views and opinions expressed are however those of the author(s) only and do not necessarily reflect those of the European Union or the European Research Council Executive Agency. Neither the European Union nor the granting authority can be held responsible for them. We are grateful to A. Stuckert, N. Brajuka, J. Sproul, and E. Tescari for their advice on repeat modelling, and J.T. Li for providing suggestions on synteny analyses. We thank M. Moore for exploratory repeat analyses.

## DATA ACCESSIBILITY AND BENEFIT-SHARING

### Data Accessibility Statement

#### Genetic data

All the genomes used in this study are available on the NCBI Genomes database (https://www.ncbi.nlm.nih.gov/genome/).

#### Code

All original code has been deposited on GitHub and is publicly available at (https://doi.org/10.5281/zenodo.7679280).

## AUTHOR CONTRIBUTIONS

Conceptualization, T.A.K., A.J.C., L.A.O., A.R., and K.C.W.V; methodology, T.A.K, N.D.Y, R.L.M, and A.R.; formal analysis, T.A.K., M.L.P.; investigation, T.A.K., N.D.Y, R.L.M., K.C.W.V, M.L.P.; resources, L.A.0. and A.R.; writing – original draft, T.A.K.; writing – review & editing, all authors; project administration, T.A.K; funding acquisition, T.A.K. and L.F.S.

