## Supplementary material for "Comparative analysis of amphibian genomes: an emerging resource for basic and applied research": Table 1

**Running title:** Comparative analysis of amphibian genomes

Tiffany A. Kosch^1^, Andrew J. Crawford^2^, Rachel Lockridge Mueller^3^, Katharina C. Wollenberg Valero^4^, Megan L. Power^4^, Ariel Rodríguez ^5^, Lauren A. O’Connell^6^, Neil D. Young^1^, and Lee F. Skerratt^1^

^1^ Faculty of Science, University of Melbourne, Melbourne, Australia

^2^ Departamento de Ciencias Biológicas, Universidad de los Andes, Bogotá, Colombia

^3^ Department of Biology, Colorado State University, Colorado, USA

^4^ School of Biology and Environmental Science, University College Dublin, Dublin, Ireland

^5^ Institute of Zoology, University of Veterinary Medicine of Hannover, Hannover, Germany

^6^ Department of Biology, Stanford University, California, USA

**Supplementary Information**

### SUPPLEMENTARY METHODS

#### Data compilation

Occurrence data were obtained for the 51 amphibian species from the Global Biodiversity Information Facility (https://www.gbif/org/; last accessed February 2024) using the R package *rgbif* (Chamberlain et al., 2022; see Table S5 for DOI's for each occurrence dataset). To ensure data quality, various error-correction steps were conducted. These steps involved removing duplicate records and incomplete coordinate data, as well as filtering out records with a coordinate uncertainty greater than 5km. Only confirmed presence records were retained for further analysis. Following the approach by Barrow et al. (2021), species with large occurrence datasets and to remove duplicates (Rhinella marina, Xenopus laevis, and *Lithobates catesbeianus*) were randomly sampled, resulting in 3000 occurrence points. For the remaining occurrence records, we extracted 19 bioclimatic metrics and elevation (m) from the WorldClim 2.0 data set (spatial resolution: 2.5 arc-min) covering the time-period 1970-2000 (Fick & Hijmans, 2017). These metrics encompass annual trends, seasonality, and extreme environmental factors, providing a more biologically meaningful representation compared to individual temperature and precipitation measurements (Biber et al., 2023) (see Table S5 for variable descriptions). We focused on temperature variables (Bio1-Bio11) given increasing evidence that TE activity is influenced by temperature (Bodelón et al., 2023). Elevation was also included which can be a proxy variable for climatic turnover and other abiotic factors such as UV radiation. To reduce overfitting of phylogenetic models, highly correlated variables were removed based on their variance inflation factor (VIF) which measures the extent to which each predictor can be explained by the other predictors (Naimi & Araújo, 2016). Using the R package *usdm*, *vifstep* was used to exclude variables with VIF values more than five. This resulted in five predictor variables; Bio2, Bio4, Bio8, Bio10 and elevation (see Table S6 for descriptions).

We summarized the genome repeats from each of the 51 amphibian species in relation to genome size resulting in three groups: proportion of total transposable elements (TEs), proportion of retroelements and proportion of DNA transposons. As a considerable proportion of repeats were unclassified, we removed these from our calculations.

#### Statistical analyses

We examined the effects of environmental variables (Bio2, Bio4, Bio8, Bio10 and elevation) on three transposable element (TE) groups, using Bayesian mixed effect models implemented in the R package *MCMCglmm* v2.34 (Hadfield, 2010). Body size was included as a covariate to account for variation in body size. Body size measures (maximum body size in mm) were obtained from the AmphiBio database (Oliveira et al., 2017), with values from Anura representing snout to vent length (SVL) and total body length (TL) for Gymnophiona and Caudata.

To enhance interpretability, separate models were constructed for each TE group and environmental traits. We obtained the phylogenetic relationships from Jetz and Pyron (2018) and trimmed the tree to include the 51 species of interest (Figure S6). To address potential phylogenetic non-independence, the trimmed phylogenetic tree was incorporated as a random factor in all models. Weakly informative priors (V = 1, nu = 0.02) were assigned to the random variance components, assuming a Gaussian error distribution. Models were run for 500,000 iterations, a thinning interval of 100 and burn-in of 10% (50,000 iterations). Parameter estimates are expressed as posterior means and 95% highest posterior density (HPD) intervals. Model convergence was assessed by examining autocorrelation values (difference between chains were all < 0.1), Heidelberg & Welch and Geweke tests were passed and ensuring effective sample sizes (ESS) for fixed and variance components were ≥ 1000 (Hadfield, 2010). The phylogenetic signal, representing the proportion of variance attributable to the phylogeny, was calculated following de Villemereuil and Nakagawa (2014) as: λ = V_PHYLO_/(V_PHYLO_ + V_RESID_). Log-transformations were applied to all three TE groups and fixed effect predictors and all were mean centered. In addition, we ran the total TE model with the removal of the three globally distributed species (*Rhinella marina, Xenopus laevis, and Lithobates catesbeianus*) to test for any potential artefacts associated with summarizing bioclimatic variables across widely distributed species.

**Supplementary Tables and Figures**

See Excel Spreadsheet for Tables S1-S4, S10-S11.

**Table S5**. Species occurrence data DOI’s from the 51 amphibian species obtained from the Global Biodiversity Information Facility (https://www.gbif/org/; last accessed February 2024).

| **Species** | **DOI** | **Species** | **DOI** |
| --- | --- | --- | --- |
| *Allobates femoralis* | https://doi.org/10.15468/dl.gx23hy | *Oophaga sylvatica* | https://doi.org/10.15468/dl.9rdzs6 |
| *Ambystoma mexicanum* | https://doi.org/10.15468/dl.5re8wr | *Pelobates cultripes* | https://doi.org/10.15468/dl.9rdzs6 |
| *Bombina bombina* | https://doi.org/10.15468/dl.f4263x | *Phrynoglossus myanhessei* | https://doi.org/10.15468/dl.jfv8pa |
| *Bombina variegata* | https://doi.org/10.15468/dl.edzsxf | *Phyllomedusa bahiana* | https://doi.org/10.15468/dl.bk55hb |
| *Bufo bufo* | https://doi.org/10.15468/dl.jnserr | *Pipa carvalhoi* | https://doi.org/10.15468/dl.52p54j |
| *Bufo gargarizans* | https://doi.org/10.15468/dl.x2t23q | *Pipa parva* | https://doi.org/10.15468/dl.7guyb3 |
| *Dendropsophus ebraccatus* | https://doi.org/10.15468/dl.q6vmg2 | *Platyplectrum ornatum* | https://doi.org/10.15468/dl.vdqgcw |
| *Discoglossus pictus* | https://doi.org/10.15468/dl.3e2kn8 | *Pleurodeles waltl* | https://doi.org/10.15468/dl.t74mnp |
| *Eleutherodactylus coqui* | https://doi.org/10.15468/dl.b33qxy | *Pseudophryne corroboree* | https://doi.org/10.15468/dl.gasa2r |
| *Engystomops pustulosus* | https://doi.org/10.15468/dl.8eued8 | *Pyxicephalus adspersus* | https://doi.org/10.15468/dl.9wucd6 |
| *Gastrophryne carolinensis* | https://doi.org/10.15468/dl.brheff | *Rana kukunoris* | https://doi.org/10.15468/dl.ws7ws9 |
| *Geotrypetes seraphini* | https://doi.org/10.15468/dl.r26aa2 | *Rana muscosa* | https://doi.org/10.15468/dl.abqykr |
| *Glandirana rugosa* | https://doi.org/10.15468/dl.gx82tt | *Rana temporaria* | https://doi.org/10.15468/dl.hfqzar |
| *Hyalinobatrachium fleischmanni* | https://doi.org/10.15468/dl.2awctk | *Ranitomeya imitator* | https://doi.org/10.15468/dl.6gahjb |
| *Hyla sarda* | https://doi.org/10.15468/dl.a2rhsd | *Rhinatrema bivittatum* | https://doi.org/10.15468/dl.gtpvkd |
| *Hymenochirus boettgeri* | https://doi.org/10.15468/dl.qv5ppp | *Rhinella marina* | https://doi.org/10.15468/dl.3k48v4 |
| *Leptobrachium ailaonicum* | https://doi.org/10.15468/dl.wgygse | *Scaphiopus couchii* | https://doi.org/10.15468/dl.485jc |
| *Leptobrachium leishanense* | https://doi.org/10.15468/dl.ksq7qa | *Scaphiopus holbrookii* | https://doi.org/10.15468/dl.a6d6sg |
| *Leptodactylus fallax* | https://doi.org/10.15468/dl.38ckf6 | *Spea bombifrons* | https://doi.org/10.15468/dl.qs5pac |
| *Limnodynastes dumerilii* | https://doi.org/10.15468/dl.f4uk7h | *Spea hammondii* | https://doi.org/10.15468/dl.83tetd |
| *Lithobates catesbeiana* | https://doi.org/10.15468/dl.f27vp2 | *Spea multiplicata* | https://doi.org/10.15468/dl.uw4g9u |
| *Lithobates sylvaticus* | https://doi.org/10.15468/dl.7fgqu8 | *Staurois parvus* | https://doi.org/10.15468/dl.w5yvkp |
| *Mantella_aurantiaca* | https://doi.org/10.15468/dl.k7ggxb | *Xenopus borealis* | https://doi.org/10.15468/dl.qzuwmt |
| *Microcaecilia_unicolor* | https://doi.org/10.15468/dl.k7ggxb | *Xenopus laevis* | https://doi.org/10.15468/dl.ckgkkj |
| *Nanorana_parkeri* | https://doi.org/10.15468/dl.fa7h4e | *Xenopus tropicalis* | https://doi.org/10.15468/dl.2aaz4d |

**Table S6.** List of Bioclimatic variables obtained from WorldClim 2.0 (<https://www.worldclim.org/>).

| **Bioclimatic variable abbreviation** | **Description** | **Units** |
| --- | --- | --- |
| Bio1 | Annual mean temperature | °C |
| Bio2 | Mean diurnal range | °C |
| Bio3 | Isothermality | % |
| Bio4 | Temperature seasonality | °C |
| Bio5 | Max temperature of warmest month | °C |
| Bio6 | Min temperature of coldest month | °C |
| Bio7 | Temperature annual range | °C |
| Bio8 | Mean temperature of wettest quarter | °C |
| Bio9 | Mean temperature of driest quarter | °C |
| Bio10 | Mean temperature of warmest quarter | °C |
| Bio11 | Mean temperature of coldest quarter | °C |
| Bio12 | Annual precipitation | mm |
| Bio13 | Precipitation of wettest month | mm |
| Bio14 | Precipitation of driest month | mm |
| Bio15 | Precipitation seasonality | % |
| Bio16 | Precipitation of wettest quarter | mm |
| Bio17 | Precipitation of driest quarter | mm |
| Bio18 | Precipitation of warmest quarter | mm |
| Bio19 | Precipitation of coldest quarter | mm |

**Table S7.** Results of Bayesian phylogenetic mixed model analyses on proportion of total transposable elements (TE’s) per genome for 51 amphibian genomes. Models predicted the proportions of total TE (log transformed) by environmental variables and body size while controlling for phylogenetic relationships. Variables have been log-transformed and mean centered. Significant effects are highlighted in bold.

| **Total TE (Body size corrected and log transformed)** | **Posterior mean** | **Lower 95% HPD** | **Upper 95% HPD** | | | |
| --- | --- | --- | --- | --- | --- | --- |
| **Variance component** |  |  |  | **λ** | **Lower 95% HPD** | **Upper 95% HPD** |
| Species (linked to phylogeny) | 0.006 | 0.002 | 0.011 | 0.555 | 0.300 | 0.786 |
| Residual | 0.004 | 0.002 | 0.007 | | | |
| **Fixed effects** |  |  |  | **pMCMC** | | |
| Intercept | 0.662 | -0.047 | 1.464 | 0.078 | | |
| Body Size | 0.013 | -0.029 | 0.051 | 0.516 | | |
| Bio2 | -0.111 | -0.269 | 0.051 | 0.156 | | |
| Bio4 | -0.008 | -0.042 | 0.024 | 0.649 | | |
| Bio8 | **-0.105** | **-0.182** | **-0.020** | **0.011** | | |
| Bio10 | 0.037 | -0.155 | 0.237 | 0.701 | | |
| Elevation | 0.003 | -0.033 | 0.043 | 0.736 | | |

**Table S8.** Results of Bayesian phylogenetic mixed model analyses on proportion of total transposable elements (TE’s) per genome (log transformed), with the removal the three invasive frog species, by environmental variables and body size while controlling for phylogenetic relationships. Variables have been log-transformed and mean centered. Significant effects are highlighted in bold.

| **Total TE (Body size corrected and log transformed)** | **Posterior mean** | **Lower 95% HPD** | **Upper 95% HPD** | | | |
| --- | --- | --- | --- | --- | --- | --- |
| **Variance component** |  |  |  | **λ** | **Lower 95% HPD** | **Upper 95% HPD** |
| Species (linked to phylogeny) | 0.006 | 0.002 | 0.011 | 0.548 | 0.298 | 0.788 |
| Residual | 0.05 | 0.002 | 0.008 | | | |
| **Fixed effects** |  |  |  | **pMCMC** | | |
| Intercept | 0.688 | -0.145 | 1.513 | 0.092 | | |
| Body Size | 0.011 | -0.035 | 0.056 | 0.623 | | |
| Bio2 | -0.113 | -0.281 | 0.058 | 0.190 | | |
| Bio4 | -0.009 | -0.046 | 0.027 | 0.635 | | |
| **Bio8** | **-0.106** | **-0.189** | **-0.021** | **0.014** | | |
| Bio10 | 0.034 | -0.165 | 0.257 | 0.742 | | |
| Elevation | 0.007 | -0.036 | 0.044 | 0.724 | | |

**Table S9.** Results of Bayesian phylogenetic mixed model analyses on proportion of total retroelements per genome. Models predicted the proportions of total retroelements (log transformed) by environmental variables and body size while controlling for phylogenetic relationships. Significant effects are highlighted in bold.

| **Total Retroelements (Body size corrected and log transformed)** | **Posterior mean** | **Lower 95% HPD** | **Upper 95% HPD** | | | |
| --- | --- | --- | --- | --- | --- | --- |
| **Variance component** |  |  |  | **λ** | **Lower 95% HPD** | **Upper 95% HPD** |
| Species (linked to phylogeny) | 0.005 | 0.002 | 0.009 | 0.616 | 0.394 | 0.827 |
| Residual | 0.003 | 0.001 | 0.005 | | | |
| **Fixed effects** |  |  |  | **pMCMC** | | |
| Intercept | 0.578 | -0.047 | 1.245 | 0.076 | | |
| Body Size | 0.008 | -0.029 | 0.042 | 0.681 | | |
| Bio2 | -0.042 | -0.182 | 0.102 | 0.558 | | |
| Bio4 | -0.011 | -0.040 | 0.019 | 0.433 | | |
| Bio8 | -0.065 | -0.136 | 0.001 | 0.061 | | |
| Bio10 | -0.012 | -0.178 | 0.152 | 0.880 | | |
| Elevation | -0.004 | -0.040 | 0.029 | 0.809 | | |

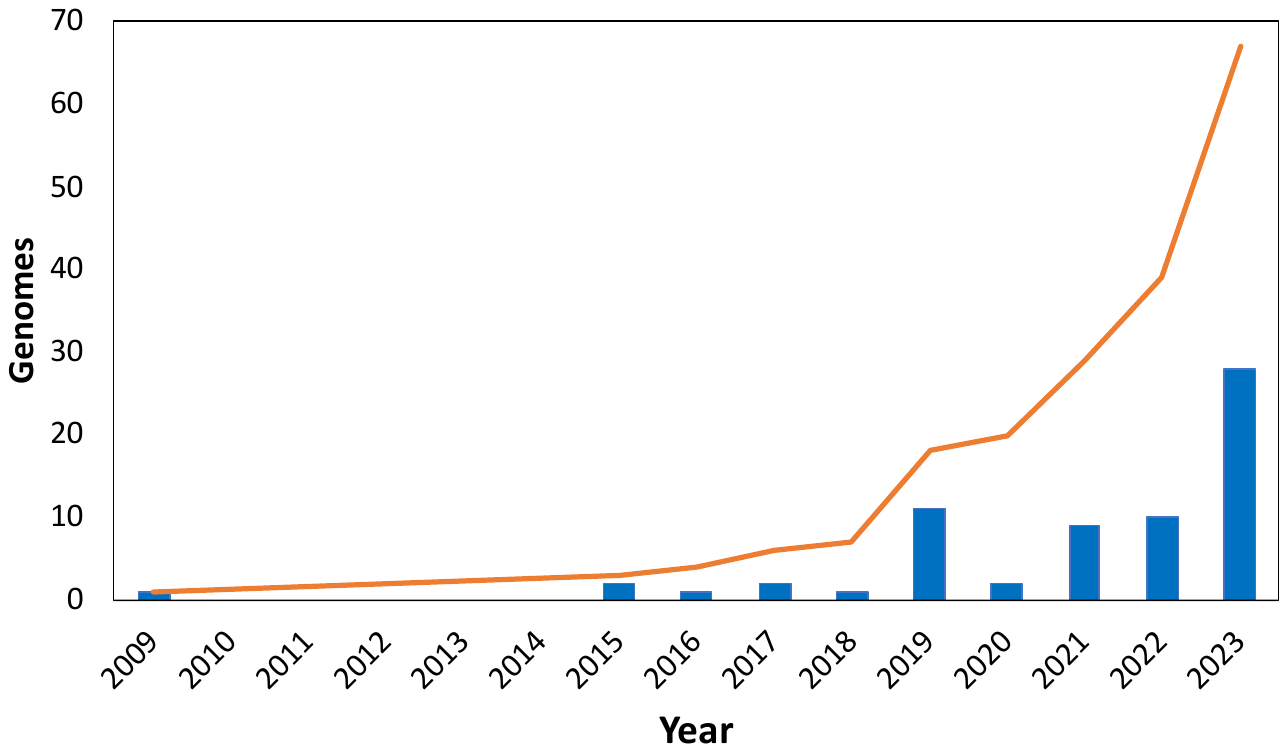

**Figure S1**. Increase in amphibian reference genomes published per year. Bars indicate counts and the line is the cumulative count.

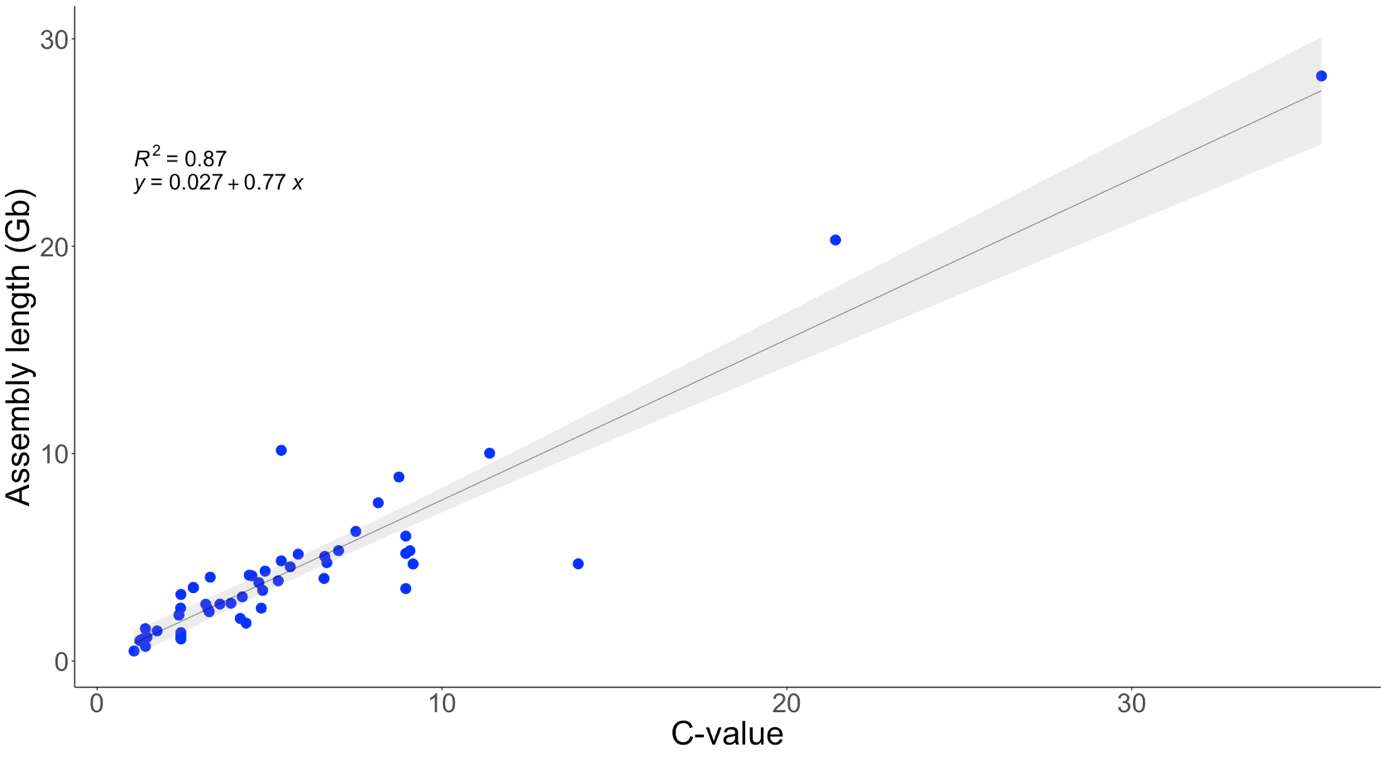

**Figure S2**. Relationship between estimated genome size (C-value) and genome assembly length. Shading represents 95% confidence interval.

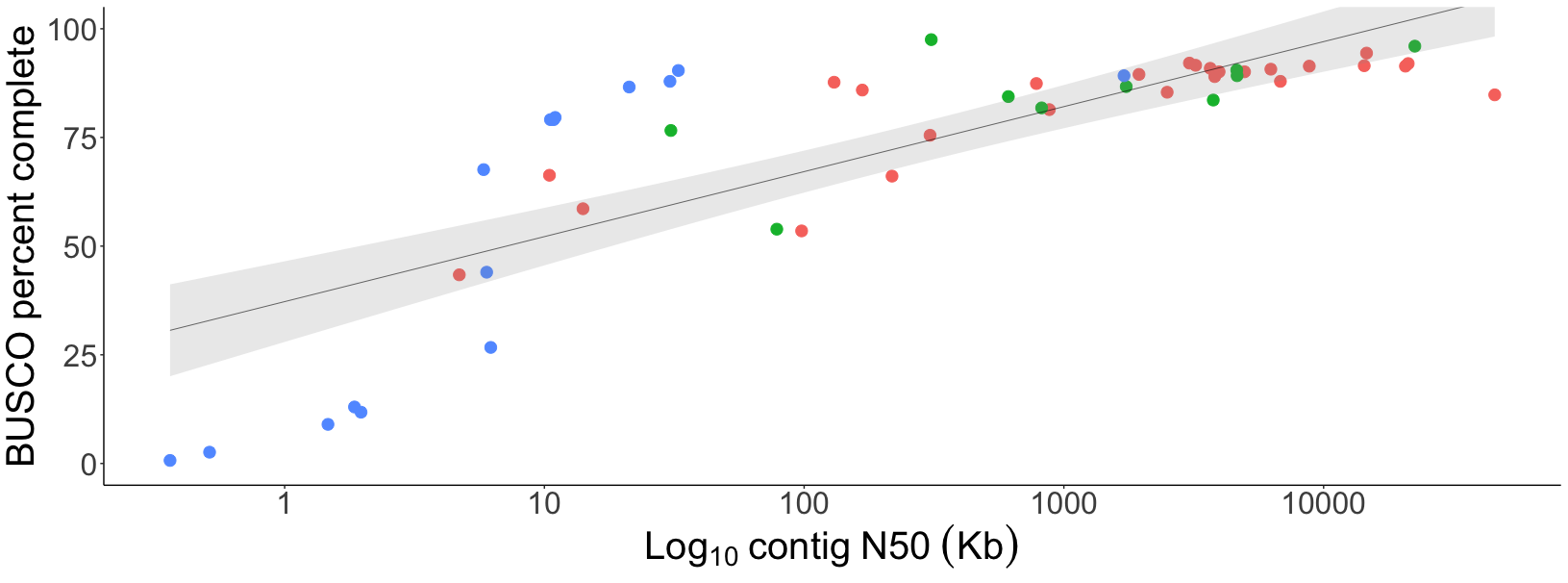

**Figure S3**. Relationship between contig N50 (genome continuity) and percentage of complete BUSCOs. (Red) hybrid assemblies, (green) long read genomes, (blue) short read genomes. Shading represents 95% confidence interval.

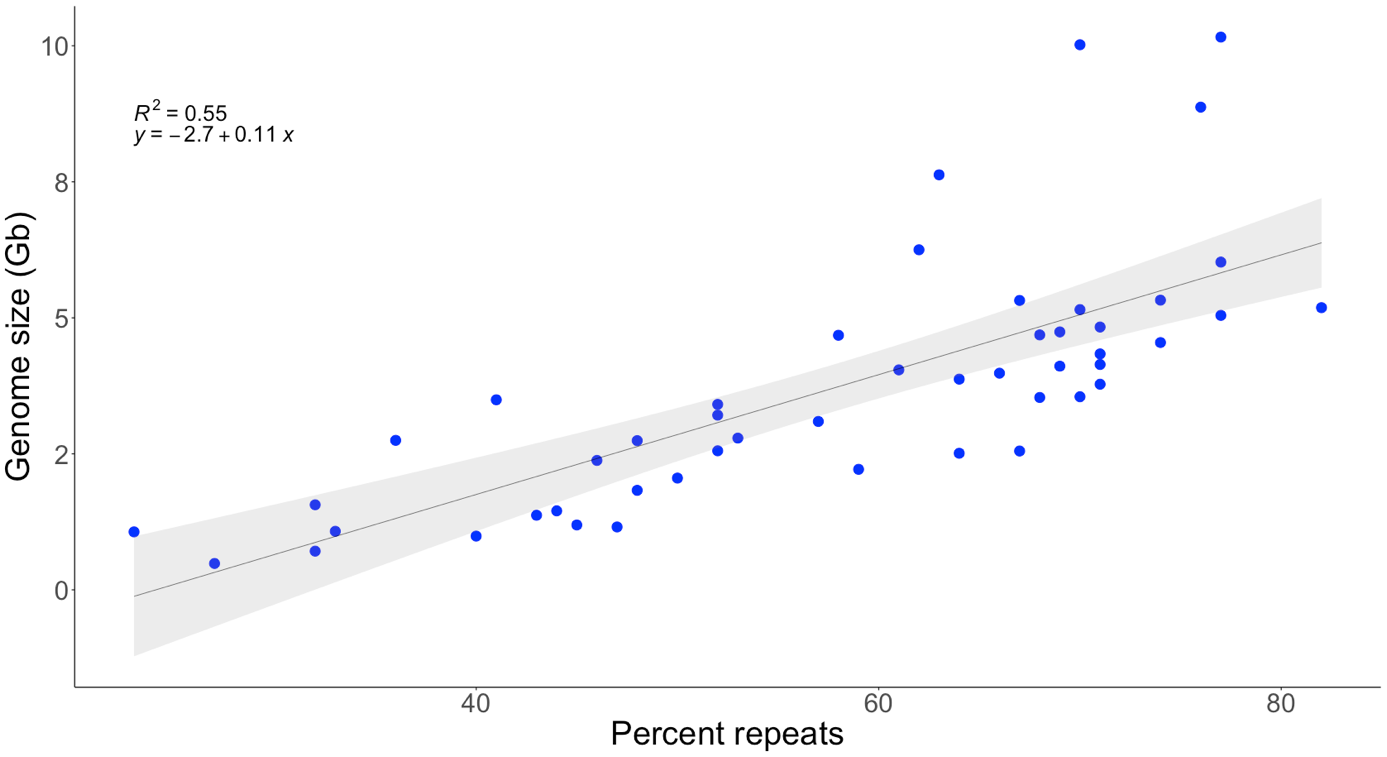

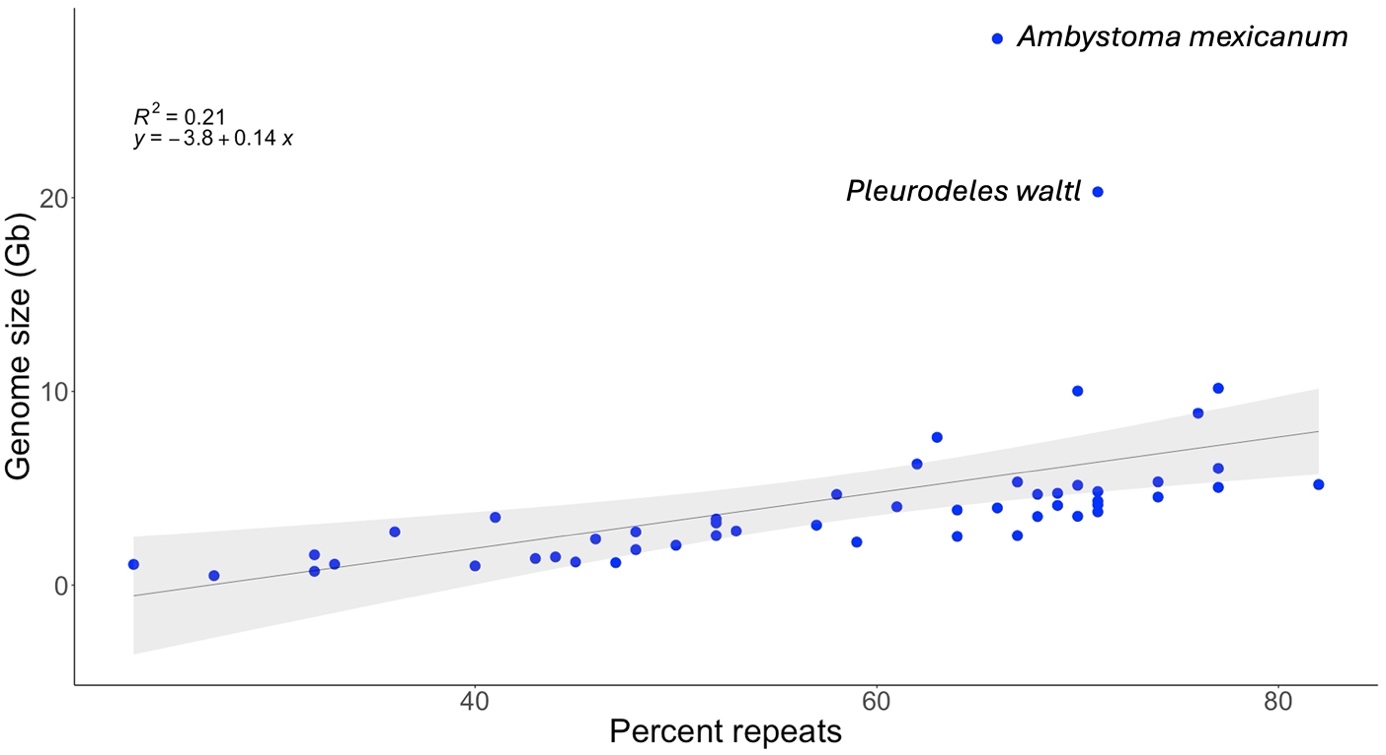

**Figure S4**. Relationship between percentage repeats and genome size. (top) Salamanders *Ambystoma mexicanum* and *Pleurodeles waltl* excluded, (bottom) *A. mexicanum and P. waltl* included. Shading represents 95% confidence interval.

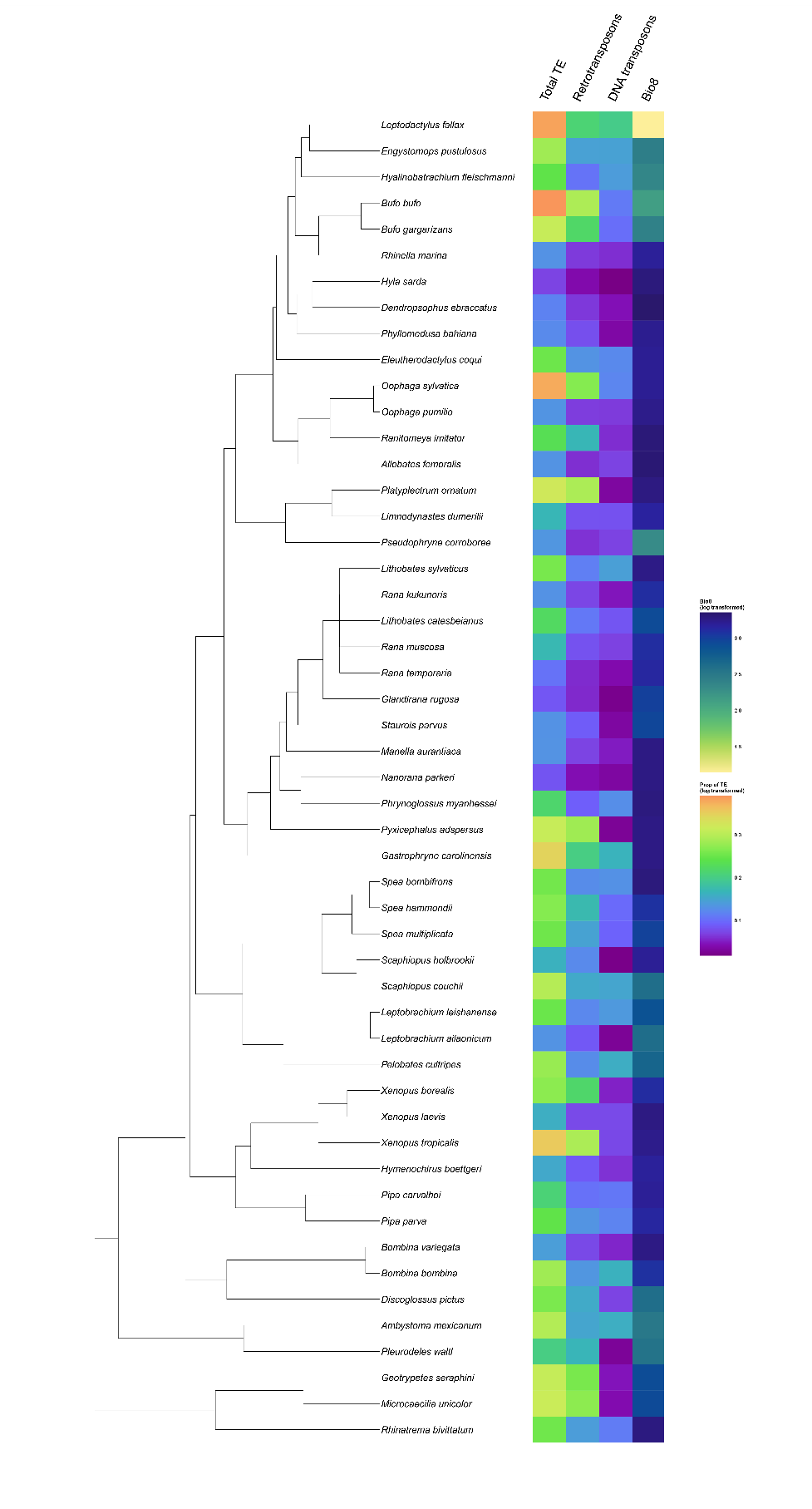

**Figure S5.** Phylogenetic heat map showing the phylogenetic distribution of transposable element (TE) groups across the 51 species and their corresponding Bio8 = mean temperature of the wettest quarter.

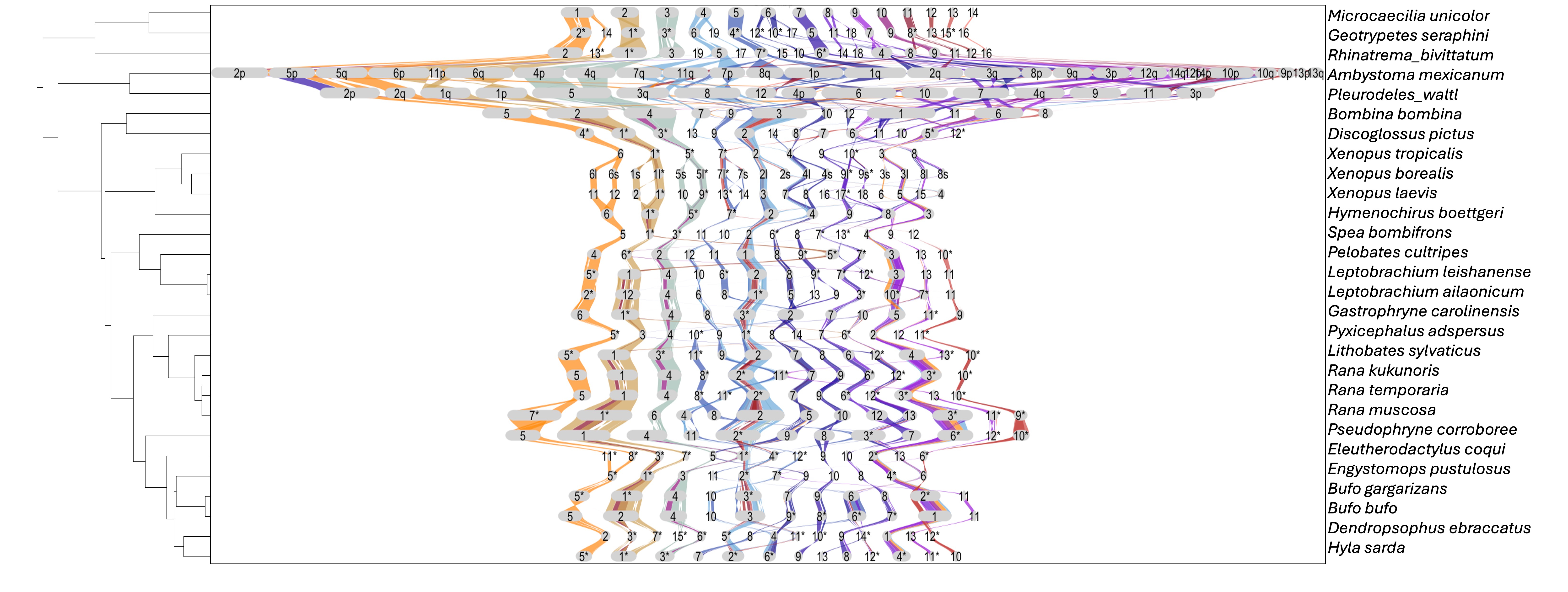

**Figure S6**. Synteny plot of BUSCOs (Benchmarking Universal Single-Copy Orthologs) for all chromosome-level amphibian genomes (N=38). Chromosomes are scaled by number of BUSCOs. Phylogenetic tree created with Timetree.org. Reference genome is *Microcaecilia unicolor*. *Indicate inverted chromosomes.

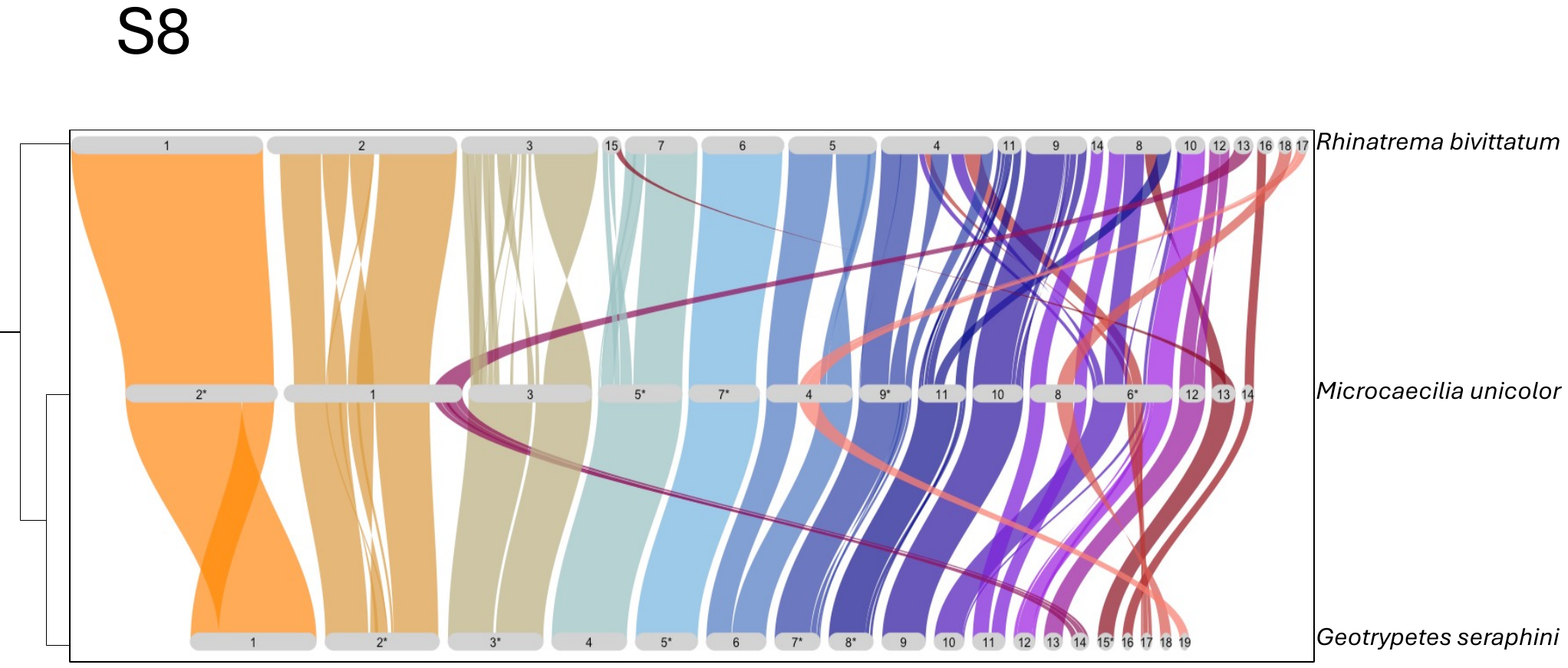

**Figure S7**. Synteny plot of BUSCOs (Benchmarking Universal Single-Copy Orthologs) for three caecilians. Chromosomes are scaled by number of BUSCOs. Phylogenetic tree created with Timetree.org. Reference genome is *Rhinatrema bivittatum*. *Indicates inverted chromosomes.

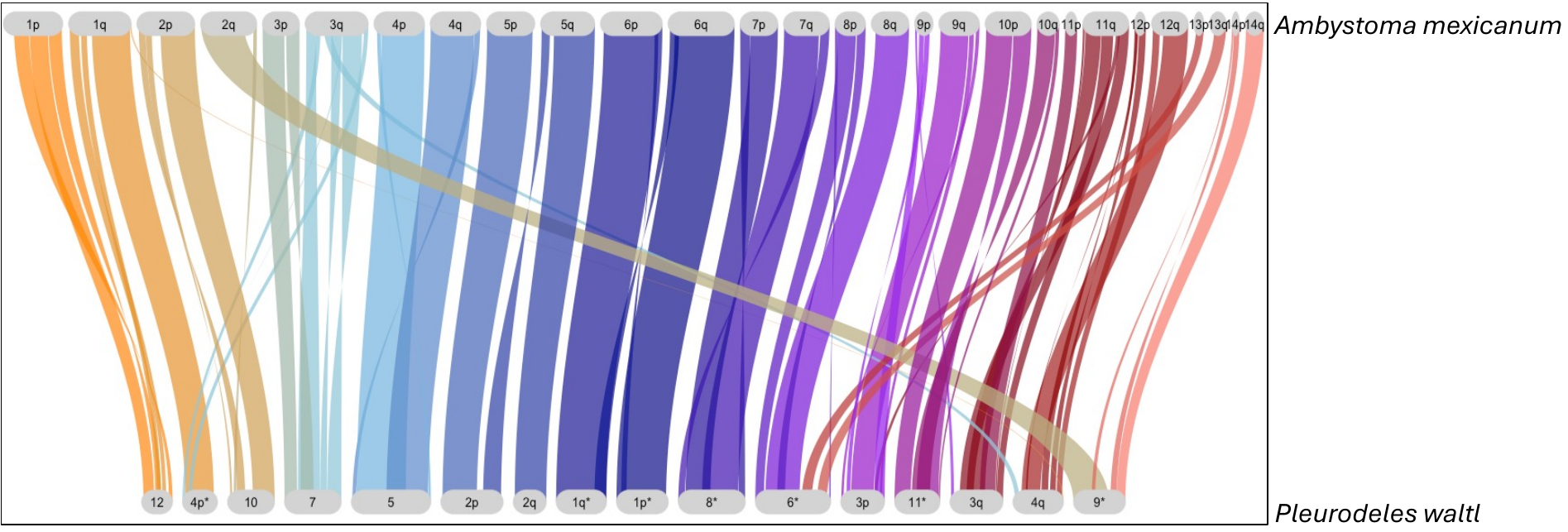

**Figure S8**. Synteny plot of BUSCOs (Benchmarking Universal Single-Copy Orthologs) for two salamanders. Chromosomes are scaled by number of BUSCOs. Phylogenetic tree created with Timetree.org. Reference genome is *Amybstoma mexicanum*. *Indicates inverted chromosomes.

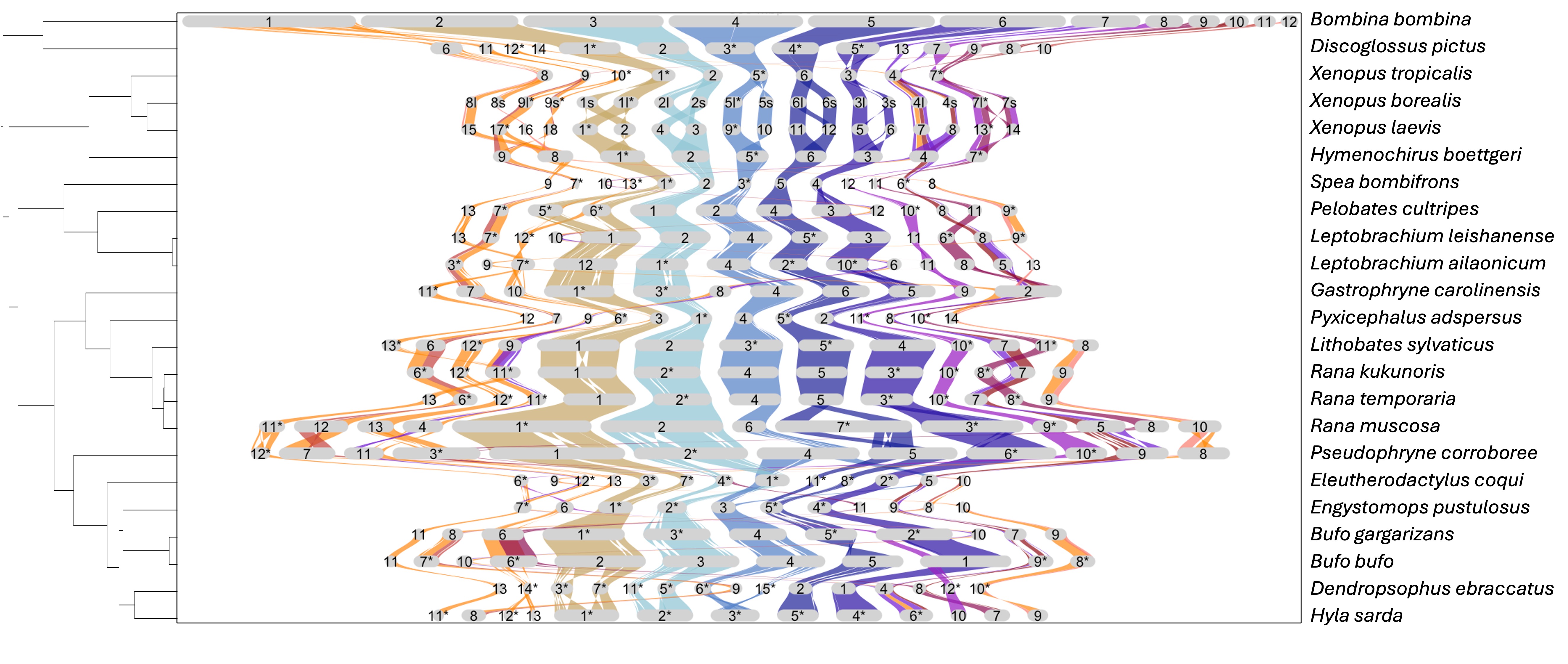

**Figure S9**. Synteny plot of BUSCOs (Benchmarking Universal Single-Copy Orthologs) for 23 frogs. Chromosomes are scaled by number of BUSCOs. Phylogenetic tree created with Timetree.org. Reference genome is *Bombina bombina*. *Indicates inverted chromosomes.
